# Moving outside the lab: markerless motion capture accurately quantifies sagittal plane kinematics during the vertical jump

**DOI:** 10.1101/2021.03.16.435503

**Authors:** John F Drazan, William T Phillips, Nidhi Seethapathi, Todd J Hullfish, Josh R Baxter

**Affiliations:** Department of Orthopedic Surgery, University of Pennsylvania, Philadelphia, Pennsylvania, United States; Electrical and Computer Engineering Department, University of Rochester, University of Rochester, Rochester, New York, United States; Department of Bioengineering, University of Pennsylvania, Philadelphia, Pennsylvania, United States

**Keywords:** biomechanics, joints angles, motion capture, mobile, DeepLabCut

## Abstract

Markerless motion capture using deep learning approaches have potential to revolutionize the field of biomechanics by allowing researchers to collect data outside of the laboratory environment, yet there remain questions regarding the accuracy and ease of use of these approaches. The purpose of this study was to apply a markerless motion capture approach to extract lower limb angles in the sagittal plane during the vertical jump and to evaluate agreement between the custom trained model and gold stand motion capture. We performed this study using a large open source data set (N=84) that included synchronized commercial video and gold standard motion capture. We split these data into a training set for model development (n=69) and test set to evaluate capture performance relative to gold standard motion capture using coefficient of multiple correlations (CMC) (n=15). We found very strong agreement between the custom trained markerless approach and marker-based motion capture within the test set across the entire movement (CMC>0.991, RMSE<3.22°), with at least strong CMC values across all trials for the hip (0.853 ± 0.23), knee (0.963 ± 0.471), and ankle (0.970 ± 0.055). The strong agreement between markerless and marker-based motion capture provides evidence that markerless motion capture is a viable tool to extend data collection to outside of the laboratory. As biomechanical research struggles with representative sampling practices, markerless motion capture has potential to transform biomechanical research away from traditional laboratory settings into venues convenient to populations that are under sampled without sacrificing measurement fidelity.

## Introduction

Marker-based motion capture is the ‘gold-standard’ technique for quantifying human movement biomechanics. However, conventional marker-based motion capture is expensive, localized to the laboratory that is inaccessible to the general population, and challenging to implement in most clinical or community settings (Lopes et al., 2018; Reinking et al., 2018). Low-cost assessment of human movement is a priority for clinicians (Parks et al., 2019). A survey of the Academy of Orthopaedic Physical Therapists in 2020 found that 47.8% of clinicians used twodimensional video analysis in patient assessment (Hensley et al., 2020), but these videos are used to provide qualitative rather than quantitative assessments of function. Similarly, there are challenges to translating biomechanical research findings into sports training and rehabilitation practice (Owoeye et al., 2020) because relying on laboratory based equipment is impractical for many practitioners (Verheul et al., 2020).

Markerless motion capture leverages computer vision techniques to address many of the limitations inherent to marker-based motion capture. One markerless motion capture tool is DeepLabCut, an open source, deep learning approach to automated pose recognition (Mathis et al., 2018). Briefly, this approach allows researchers to assemble a set of labeled training images and train a neural network to automatically track the position of user selected, manually identified landmarks throughout a video (Mathis and Mathis, 2020). This tool’s flexibility is a major strength over other markerless motion capture paradigms. Rather than tracking a predetermined list of external features or landmarks, it can track any feature of interest. It has been used in disparate applications ranging from tracking the muscle-tendon junction during muscle contractions imaged using ultrasound (Krupenevich et al., 2020) to tracking human movement outside of the laboratory in challenging conditions like underwater running (Cronin et al., 2019). However, this tool has yet to be widely used in the field of biomechanics. One likely reason for this underutilization may be lack of direct comparisons with gold standard marker-based motion tracking systems (Colyer et al., 2018).

The purpose of this study was to compare sagittal hip, knee, and ankle kinematics measured using a custom-trained markerless motion capture solution with a ‘gold standard’ marker-based motion capture system. We leveraged a publicly-available data set containing both digital video and marker-based motion capture data of 90 subjects performing a variety of activities of daily living (Ghorbani et al., 2020). We used these data to determine the effectiveness of DeepLabCut as a markerless motion capture alternative to marker-based motion capture. We expected that lower extremity sagittal kinematics during a counter movement jump would track similarly well using markerless motion capture compared to marker-based motion capture.

## Methods

### Data Source and Preparation

We trained and validated our model using a publicly available data set containing marker-based motion capture and time synchronized commercial video capture of 90 subjects (Ghorbani et al., 2020). Data were collected at 120 Hz using a 15 camera motion capture system (Qualisys 300 and 310). Two dimensional video were captured at 30 Hz using two perpendicularly placed cameras (Sony ICX285, 800×600 pixels) which corresponded to sagittal and coronal views relative to the starting subject position. Each of the 90 subjects performed 20 actions during a data collection session pertaining to activities of daily life while outfitted in standardized clothing and a 67-point marker set. We opted to evaluate the vertical jump due to its ubiquity as a functional test in athletics and physical rehabilitation. We identified starting and ending time points for the vertical jump portion of the video for each subject (Fig. 1 D). We excluded 5 of the 90 subjects from further analysis due to incomplete motion capture data or data entry errors.

**Figure 1:**
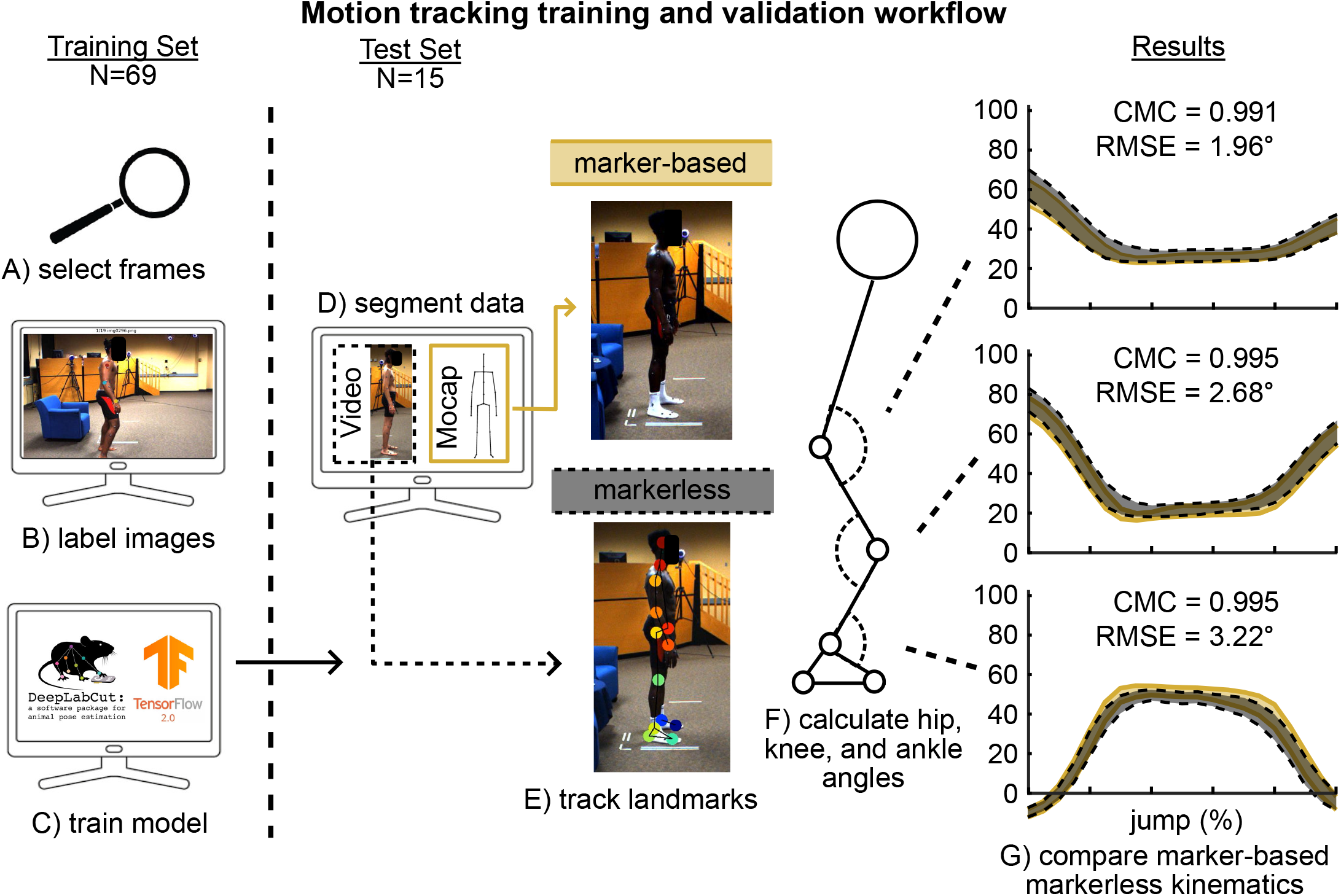
A) Six frames per video within the training set (N=69, two videos each) were selected using the automated frame selection feature of deep lab cut for labeling. B) 19 landmarks on 828 images were labeled by four different investigators using the deep lab cut provided GUI. 60 images were shared between blinded investigators to evaluate agreement in labeling. C) We trained the model on labeled images using ResNet 50 on a personal computer with a discrete graphics card. D) We segmented both video and marker-based motion capture into in the test set (N=15) into vertical jump trials. E) We applied the trained model to track 19 landmarks for each video within the test set. F) We calculated hip, knee, and ankle angles in the sagittal plane for both marker-based and markerless motion capture methods. G) We compared hip, knee, and ankle angles between marker-based (gold) and markerless (black) across all identified vertical jumps (N=25) using Coefficients of Multiple Correlations (CMC) and Root Mean Square Error (RMSE). Shaded regions bordered by dashed lines represent 95% confidence intervals.

### Development and training of markerless motion capture model

We randomly assigned our remaining subjects into two groups; the training set (69 subjects) and the testing set (16 subjects) (Fig. 1). We trained our model with labeled images from 69 subjects. To capture full body motion, we decided to define a bilateral model with 19 landmarks primarily in reference to joint centers. This label set included the head, neck, shoulders, elbows, wrists, pelvis, hips, knees, ankles, heels, and toes. For each subject, we extracted six images from each of the two cameras for labeling using an automated feature in DeepLabCut that identified distinct frames optimized for labeling, resulting in 828 labeled images for model training (Fig. 1 A). Images were grouped by subject and randomly assigned to one of four investigators for labeling (Fig. 1 B). Because activities of daily living often result in body parts being occluded, we instructed labelers to not label landmarks not clearly visible. To establish agreement between labelers, 5 random subjects (60 total images) were labeled by four blinded labelers. We evaluated degree of agreement in landmark placement between labelers using the C-1 formulation of the Intraclass correlation coefficient (McGraw and Wong, 1996; Trevethan, 2017) and found very strong agreement between these four labelers (ICC = 0.998, RMSE = 4.52 pixels) within a subset of subjects (n=5 across four raters). We then randomly selected a labeler for each of the common subjects for inclusion in the final training set.

Previous studies have described the technical aspects of model training in detail (Cronin et al., 2019; Mathis and Mathis, 2020). Therefore, we will focus on how we trained the model in practice while providing a detailed human biomechanics-focused user guide for reference in appendices. We trained the model on a personal computer with a discrete graphics card (Nvidia GeForce 2080 Super) using the built-in graphical user interface for DeepLabCut (Fig. 1 C). Training the model using the ResNet 50 neural network took approximately eight hours of computational time. Using a personal computer with a discrete graphics card is helpful for model building and tracking, but the DeepLabCut team provide fully documented guides to leveraging free computing resources like Google Colab.

Following training, we used this new model to track the 16 subjects in the test set (Fig. 1 E). We filtered all data using a 6 Hz low pass filter. We calculated hip (*θ_H_*), knee (*θ_K_*), and ankle (*θ_A_*) sagittal plane angles using marker-based and markerless motion capture using the position of joint centers and the dot product (Fig. 1 F). To account for differences in marker placement and the visually defined landmarks, we applied an angular position offset to each markerless tracking trial. We calculated this angular position offset by taking the difference between the marker-based and markerless derived joint angles with each subject standing in a neutral position. We identified and segmented the 1-3 vertical jumps that each subject performed during the trial. We excluded one subject due to poor jumping form (bouncing on toes rather than performing a vertical jump) which resulted in a test set sample of 15 subjects, with 25 jumps across all subjects.

### Evaluation of Model Performance

To assess the performance of our markerless motion capture model, we calculated the root mean square error of the entire jump movement and the coefficient of multiple correlations (CMC, Kadaba et al., 1989) between the gold standard marker-based motion capture and markerless motion capture. We calculated coefficients of multiple correlations (r) for each individual jump as well as averaged trials across the entire movement. These coefficients range from 0 to 1 where higher values represent great agreement between two measurement methods. Specifically, r values between 0 to 0.36 represent poor agreement, 0.36 to 0.67 represent moderate agreement, 0.67 to 0.9 represent strong agreement, and 0.9 to 1.0 represent very strong agreement.

## Results

We found very strong agreement between the custom trained markerless motion capture and marker-based motion capture (CMC>0.991, RMSE<3.22 °) (Fig. 1 G). Measurement techniques had strong agreement for hip sagittal motion (0.853 ± 0.23) and very strong agreement for both knee (0.963 ± 0.471) and ankle (0.970 ± 0.055) sagittal motion.

## Discussion

In this study we evaluated the performance of a custom-trained markerless motion capture model by comparing lower-extremity sagittal kinematics to a gold standard marker-based motion capture system across a large publicly available data set. We calculated hip, knee, and ankle kinematics during vertical jumps performed by 15 subjects and found strong to very strong agreement between these two measurement approaches (Fig 1. G). These findings agree with previous studies comparing alternatives to motion capture (Colyer et al., 2018). The strong agreement between marker-based and markerless motion capture provides evidence that markerless motion capture is a viable tool to collect data outside of the lab with a properly trained model.

There are several limitations to this study. We utilized markerless motion capture to measure planar kinematics and joint angles rather than combining multiple camera angles to approximate true 3-dimensional motion capture as shown by other studies (D’Antonio et al., 2020; Schwarcz and Pollard, 2018). These 3-dimensional approaches require careful camera placement as well as pre-calibration which was not possible with the preexisting data set we used. Another limitation is that we only evaluated the vertical jump rather than other activities such as walking or jogging. We selected the vertical jump due to its widespread use as a benchmark for physical testing. Additionally, we only tracked the leg that was closest to the camera due to occlusion of the rear leg and to demonstrate a simple way to collect data that extends the reach of biomechanics research. Despite these limitations, we believe our work demonstrates DeepLabCut provides researchers and clinicians with an adaptable tool to collect biomechanical data outside the lab that is comparable to motion capture.

Despite the increasing prevalence of open source tools for markerless motion capture, it is still in the early stages of adoption in the field of human biomechanics (Colyer et al., 2018). As motion capture represents significant capital investments and has been the gold standard for decades, this hesitance is understandable. We believe that markerless motion capture is most valuable as an additional tool that expands research opportunities rather than as an outright replacement for present lab based systems. Markerless motion capture empowers researchers to use consumer-grade cameras to measure human movement outside of traditional laboratory settings that were previously inaccessible to researchers (Seethapathi et al., 2019).

Reliably tracking human motion outside of the laboratory setting has potential to vertically advance biomechanical research practice. Although other fields in human subject research have grappled with and are now working to address issues of unrepresentative sampling practices (Bornstein et al., 2013), the field of biomechanics continues to struggle in this domain. A recent review of 30 years of publications within the Journal of Applied Biomechanics found that 84% of studies used convenience samples of less than 30 subjects (Vagenas et al., 2018). The widespread use of convenience sampling in biomechanics is problematic. The individuals at high risk of musculoskeletal pathologies – the elderly, minorities, and socioeconomically marginalized – are not convenient to laboratory-based researchers. Minority and low-income individuals are significantly less likely to seek office-based medical treatment for early treatment of a musculoskeletal condition relative to others (Rabah et al., 2020). Even when covered by the same medical insurance, those from socioeconomically disadvantaged neighborhoods receive less musculoskeletal care (Rahman et al., 2020). Similarly, subjects “conveniently” drawn from research staff are college educated, which affects musculoskeletal disease pathology (Barbour et al., 2017). Moreover, biomedical engineers, like many other fields in science, technology, engineering, and mathematics (STEM), are not diverse (Linsenmeier and Saterbak, 2020). Also, as convenience sampling often relies on university students, the persistent underrepresentation of Black and Hispanic students at four-year research universities further inhibits representative convenience sampling (Roy, 2019).

Increasing diversity among research participants is an unmet challenge in biomedical research writ-large (Oh et al., 2015) and situating biomedical research studies within community accessible venues has been identified as an important strategy for overcoming barriers to subject recruitment (Heller et al., 2014). Markerless motion capture has potential – like other devices including mobile dynamometry (Drazan et al., 2020), instrumented insoles (Hullfish and Baxter, 2020), tensiometers (Martin et al., 2018), and inertial measurement units (Seel et al., 2014) – to transform biomechanical research away from traditional laboratory settings into venues convenient to populations that are presently undersampled without sacrificing measurement fidelity.

## Supporting information

Simple DLC User Guide

## Acknowledgements

This work was funded in part the PennPORT IRACDA Fellowship Program (NIH grant# K12GM081259), NIH/NIAMS K01AR075877, and the Penn Center for Musculoskeletal Disorders (NIH/NIAMS P30AR069619).

