## Supplementary material for "Moving outside the lab: markerless motion capture accurately quantifies sagittal plane kinematics during the vertical jump": Simple DLC User Guide

### Overview of using DeepLabCut for human biomechanics projects

- While the creators of DeepLabCut have provided significant documentation that is supremely helpful, the sheer amount of documentation and the myriad of applications can make it difficult to navigate for first time users with simple use cases.
- We trained and tested our model on the MoVi data set which is a publically available data set containing commercial video, marker-based motion capture data, and IMU data for 90 subjects.
- **The purpose of this document is to provide a simplified walkthrough of how to use DeepLabCut for markerless human motion capture.**

### Launching DeepLabCut

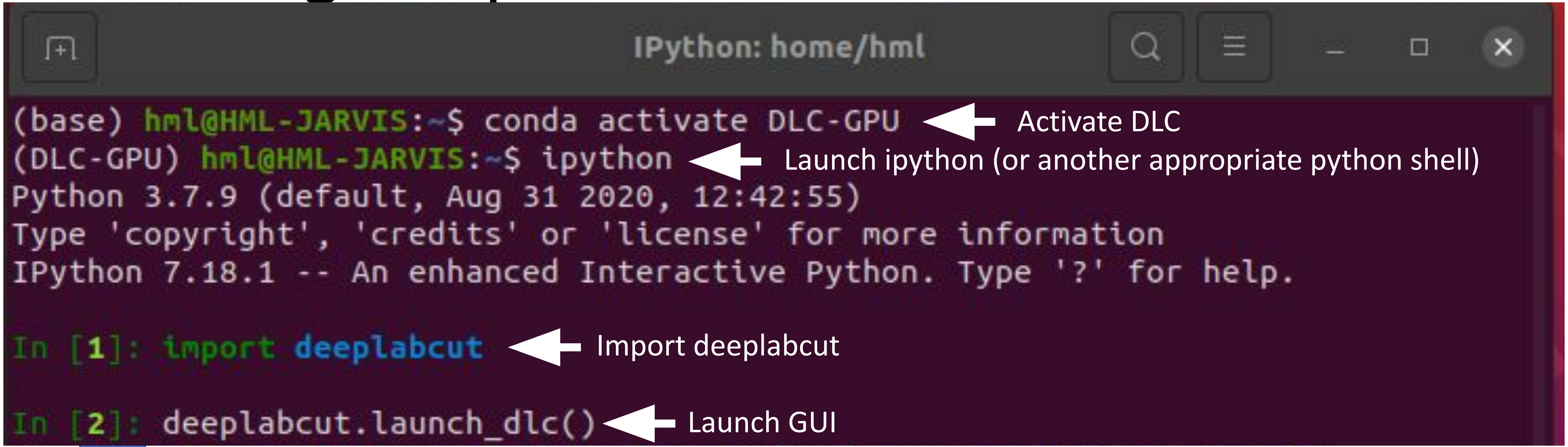


GUI Following Launch

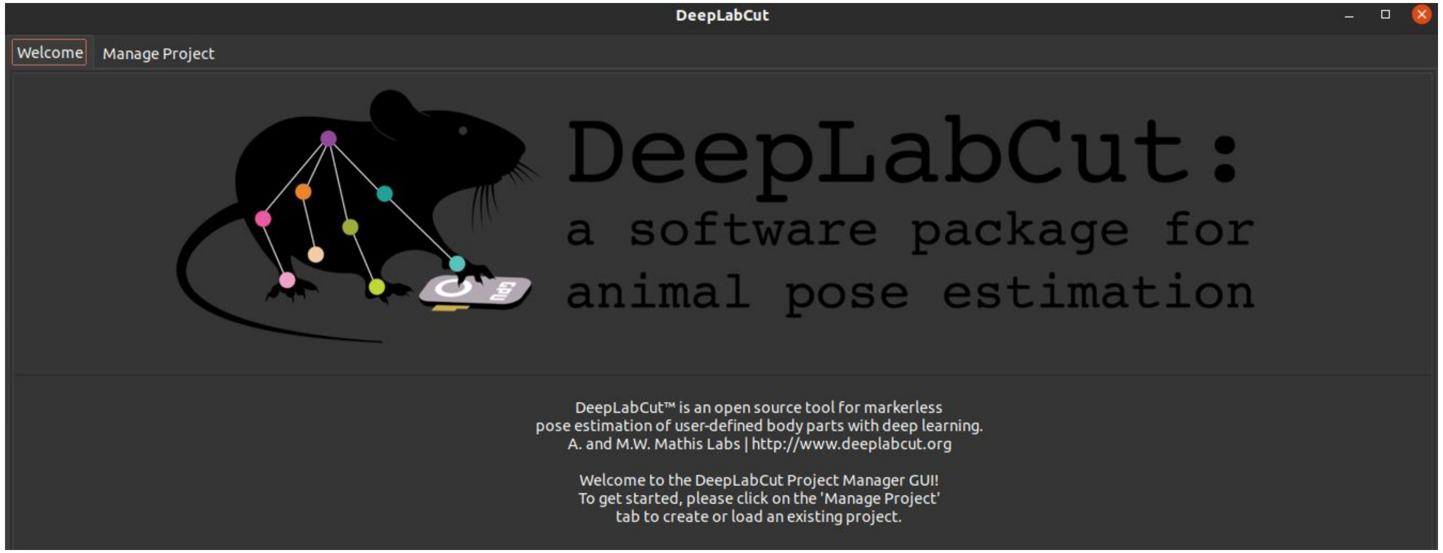

### Manage Project Tab

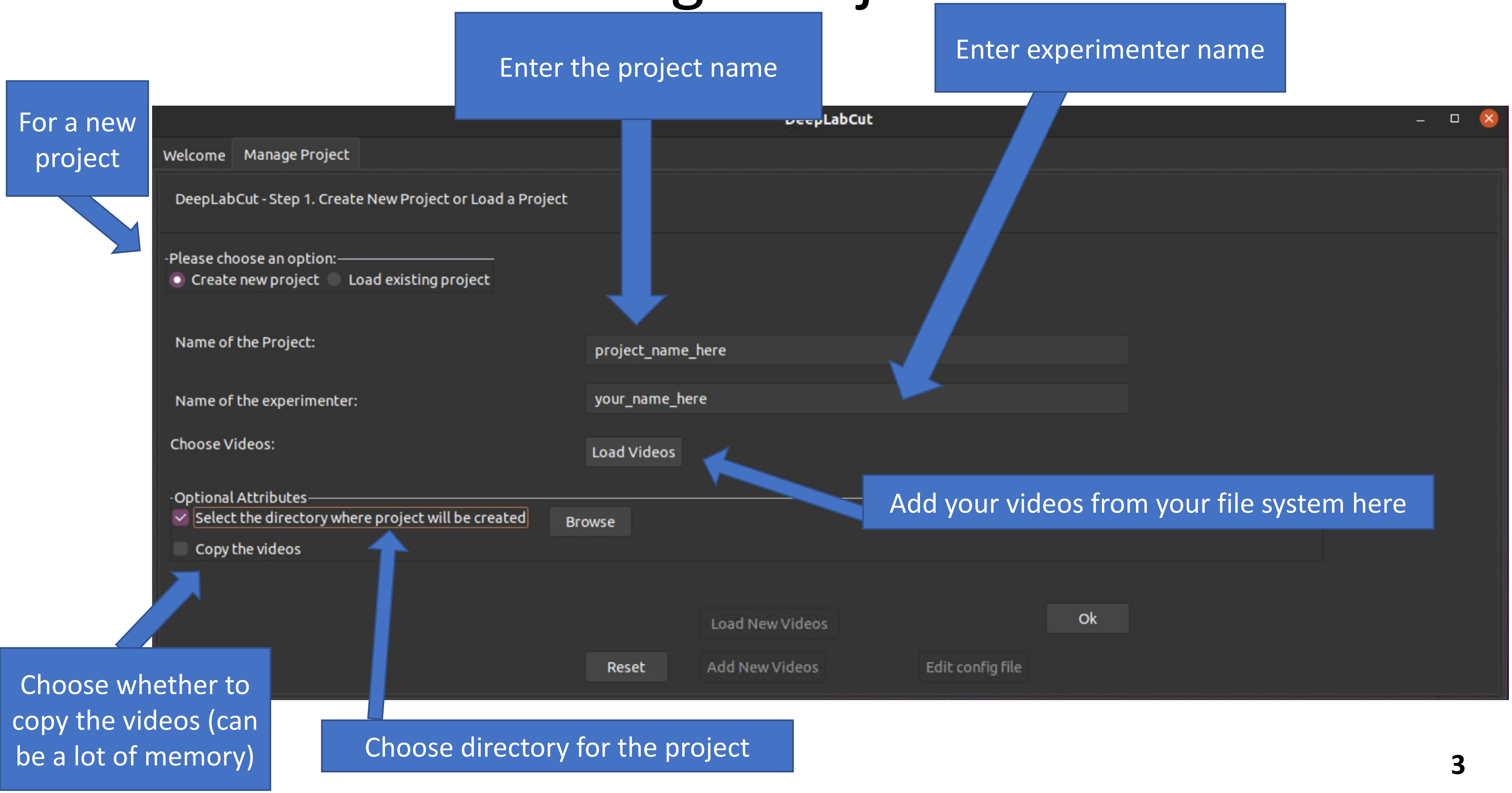

### Once a project is created/loaded, the GUI creates a series of tabs to walk you through the process:

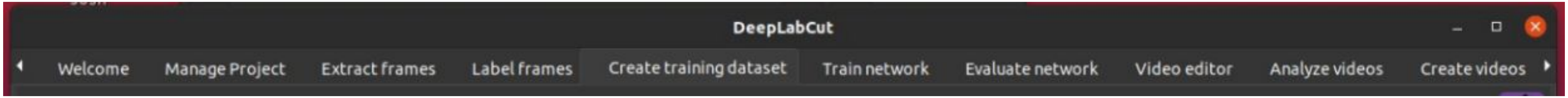


A B C D E F

A. Extracting frames for labeling
B. Labeling frames to provide training set
C. Creating training set from labeled images
D. Train the network using the training set
E. Evaluate performance of trained network
F. Analyze new videos using trained networks

This is meant to be a quick overview. For more details on every parameter, see documentation from the creators of DLC

**When creating a NEW model, start at A**

**When applying an existing model, start at F**

### A) Extracting frames

Config File Path (next page more detail)

Automatic or manual frame selection

If automated, either select based on uniform interval or by kmeans clustering

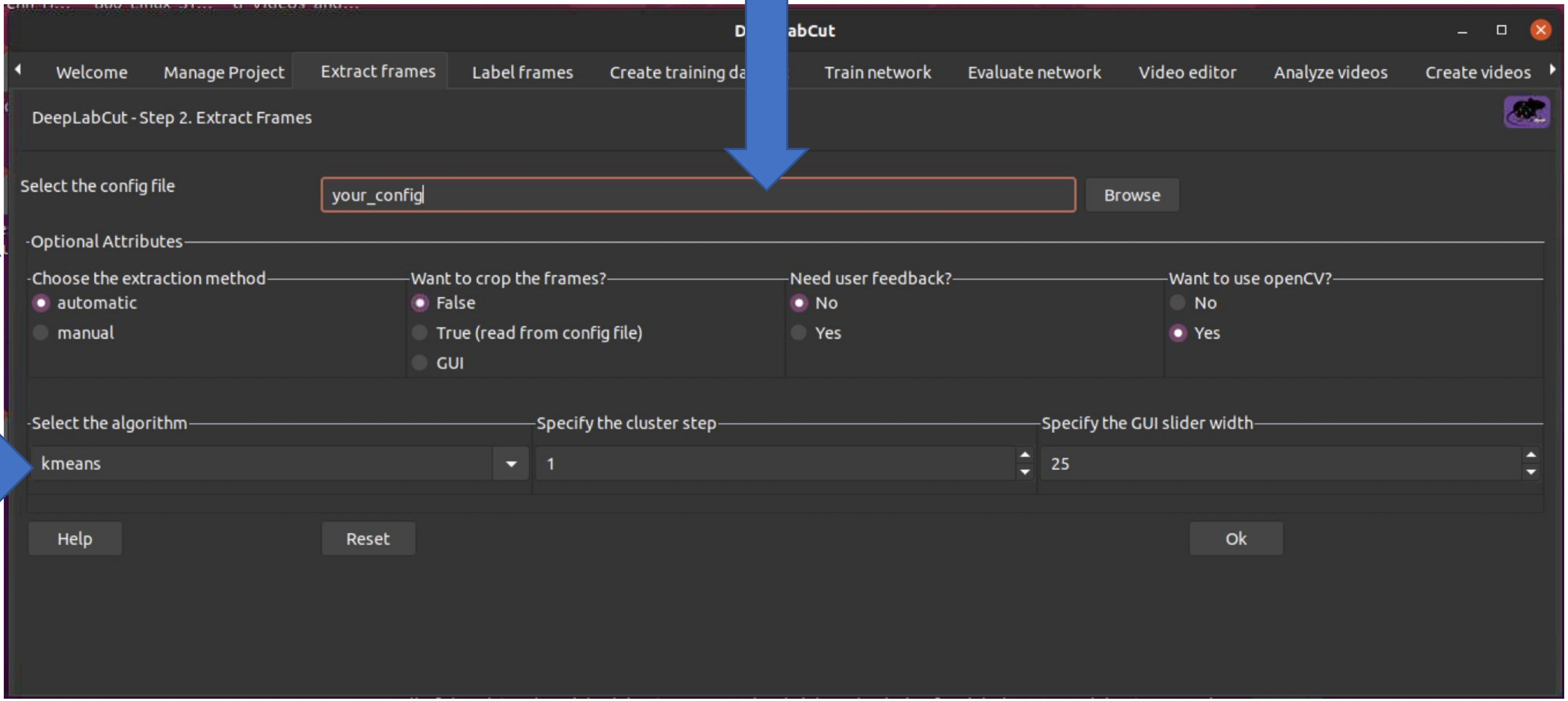

### Editing Config File

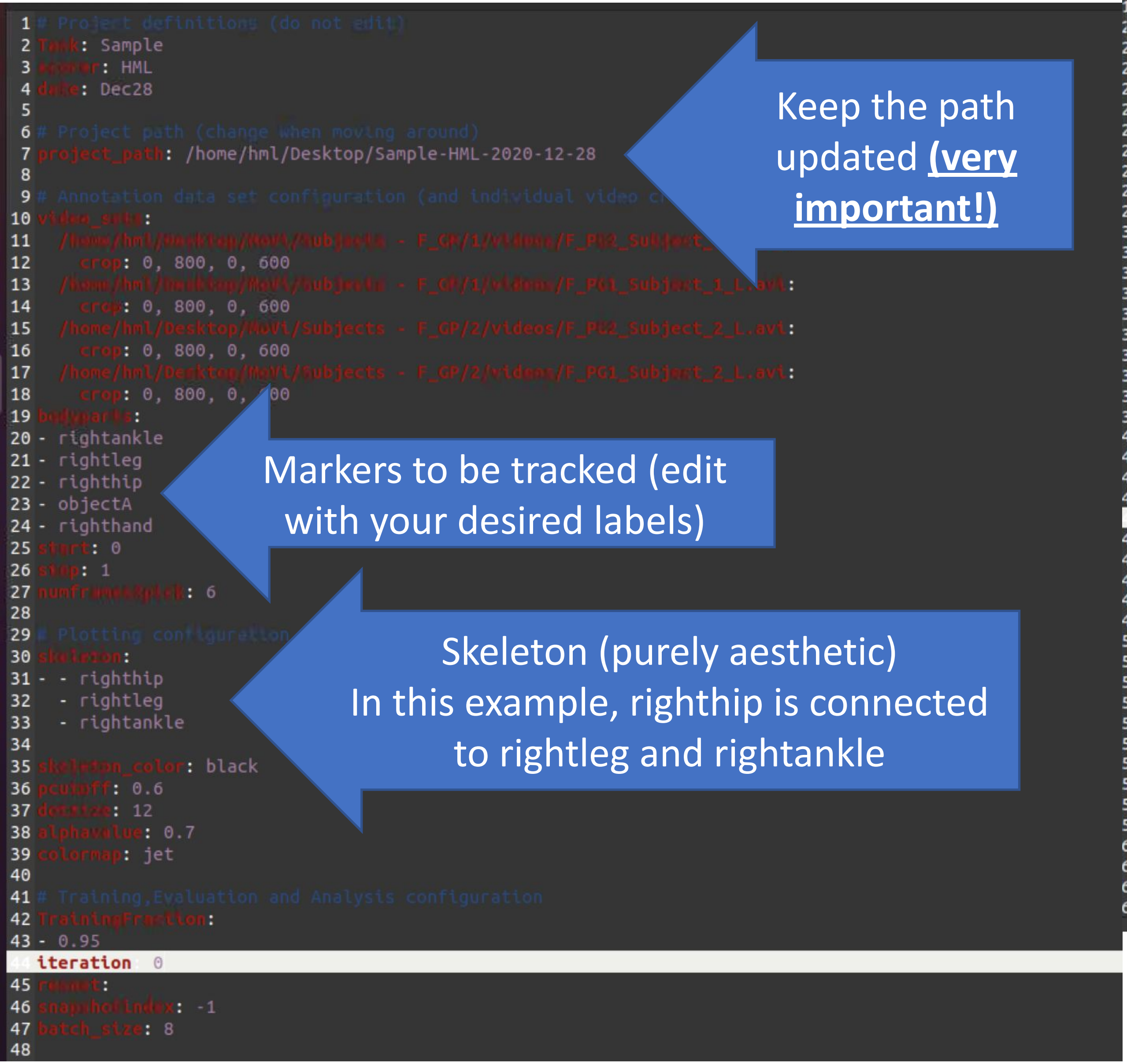


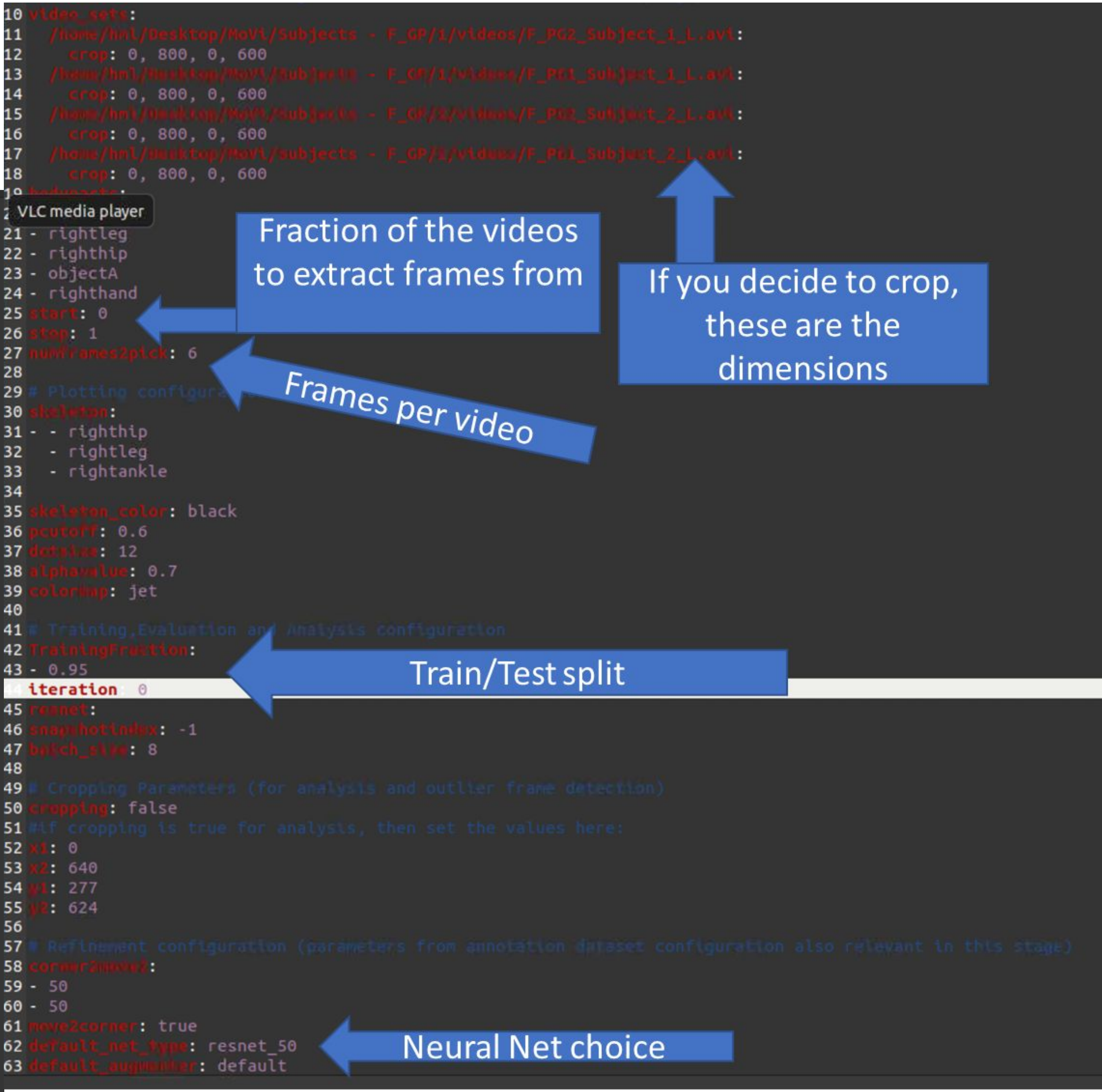

### More Config File editing!

More details on config file parameters from the creators of DeepLabCut

Source:
[https://github.com/DeepLabCut/DeepLabCut/blob/master/docs/functionDetails.md](https://github.com/DeepLabCut/DeepLabCut/blob/master/docs/functionDetails.md)

#### BOX 1: Glossary of parameters in the project configuration file (config.yaml)

*The config.yaml file sets the various parameters for generation of the training set file and evaluation of results.*
*The meaning of these parameters is defined here, as well as referenced in the relevant step.*

**Parameters set during the project creation:**

**task:** Name of the project (e.g. mouse-reaching). ( do not edit)
**scorer:** Name of the experimenter (do not edit)
**date:** Date of creation of the project. (do not edit)
**project_path:** Full path of the project; edit this if you need to move the project to a cluster/server/another computer or a different directory on your computer
**video_sets:** A dictionary with the keys as the full path of the video file and the values, crop as the cropping parameters used during frame extraction.
(use the function add_new_videos to add more videos to the project; if necessary the paths can be edited manually, and the crop values are designed to be edited manuall

**Important parameters to edit after project creation:**

**bodyparts:** List containing names of the points to be tracked. The default is set to hand, Finger1, Finger2, Joystick. Do not change after labeling frames (and saving labels).
You can *add* additional labels later, if needed.
**numframes2pick:** This is an integer that specifies the number of frames to be extracted from a video or a segment of video. The default is set to 20.
**colormap:** It specifies the colormap used for plotting the labels in images or videos in many steps.
Matplotlib colormaps are possible (https://matplotlib.org/examples/color/colormaps_reference.html).
**dotsize:** Specifies the marker size when plotting the labels in images or videos. The default is set to 12.
**alphavalue:** Specifies the transparency of the plotted labels. The default is set to 0.5.
**iteration:** This keeps the count of the number of iterations used to create the training dataset. The first iteration starts with 0 and thus the default value is set to 0.
Do not change this manually.

If you are extracting frames from long videos:
**start:** Start point of interval to sample frames from when extracting frames. Value in relative terms of video length, i.e. [start=0,stop=1] is the full video. The default is set to 0.
**stop:** Same as start, but the end of the interval. Default is 1.

**Related to the Neural Network Training:**

**TrainingFraction:** This is a two digit floating-point number in the range [0-1] to split the dataset into training and testing dataset. The default is set to 0.95.
**resnet:** This specifies which pre-trained model to use. The default is set to 50 (user can choose 50 or 101, see also Mathis et al, 2018).

**Used during video analysis:**

**batch_size:** This specifies how many frames to process at once during inference (For tuning of this parameter see Mathis & Warren 2018).
**snapshotindex:** This specifies which checkpoint to use to evaluate the network. The default is set to -1. Use *"all"* to evaluate all the checkpoints.
Snapshots refer to the stored TensorFlow configuration, which holds the weights of the feature detectors.
**p-cutoff:** This specifies the threshold of the likelihood and helps distinguishing likely body parts from uncertain ones. The default is set to 0.1.
**cropping:** Specifies if the analysis video needs to be cropped. The default is set to False.
**x1,x2,y1,y2:** These are the cropping parameters used for cropping novel video(s). The default is set to the frame size of the video.

**Used during refinement steps:**

**move2corner:** In some (rare) cases the predictions from DeepLabCut will be outside of the image (due to the location refinement shifts).
This binary parameter makes sure that those points are mapped to a user defined point within the image so that the label can be manually moved to the correct location. The default is set to True.

**corner2move2:** This is the target location, if move2corner is True. The default is set to (50,50).

### B) Labeling Frames

Place markers by right-clicking

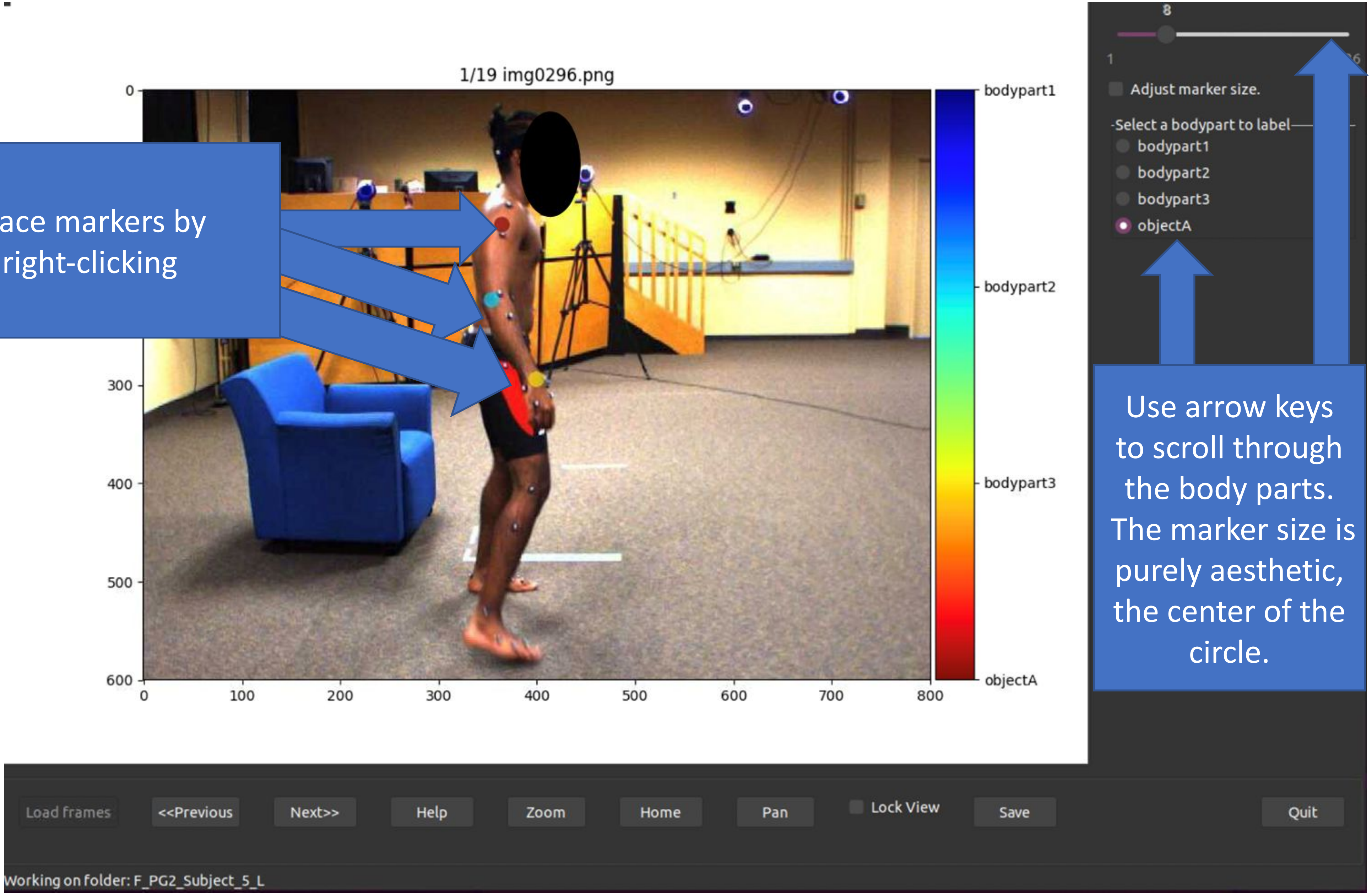


Use arrow keys to scroll through the body parts. The marker size is purely aesthetic, the center of the circle.

### C) Create Training Set

Choose your neural net

Augmentation method selection, see link at bottom of screen for more options, but default is generally good

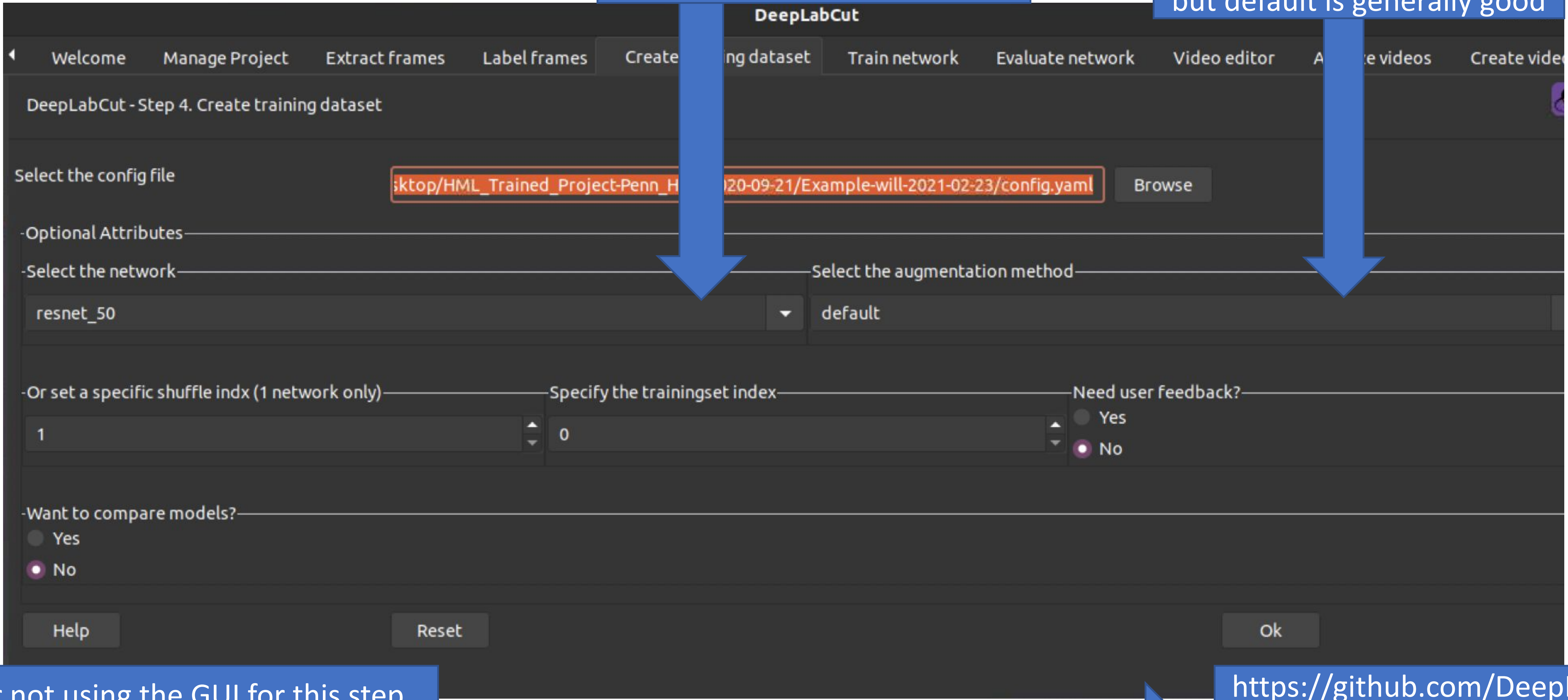


Consider not using the GUI for this step, some options (like converting Windows-labeled videos to Linux) may not be available via GUI

Ex. deeplabcut.create_training_dataset(config_path, augmenter_type='imgaug')

https://github.com/DeepLabCut/DeepLabCut/wiki/DOCSTRINGS#create_training_dataset

### D) Training the Network

May have to restrict GPU. If so, restart your python shell with this command (or a similar one import tensorflow as tf
gpu_options = tf.GPUOptions(per_process_gpu_memory_fraction=0.8)
sess = tf.Session(config=tf.ConfigProto(gpu_options=gpu_options))

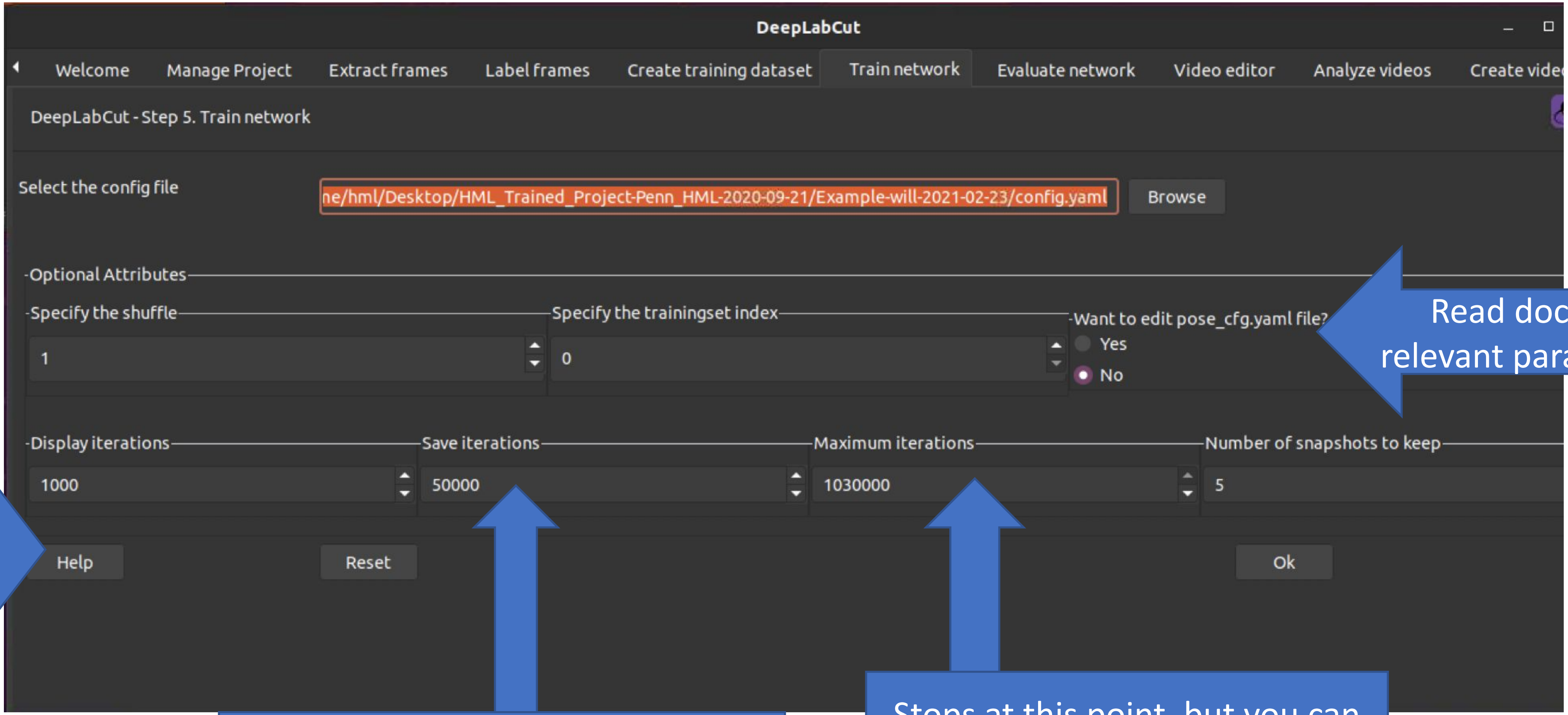


Read docs for relevant parameters

To keep track of progress

How often to backup training

Stops at this point, but you can stop the program anywhere. 200k is recommended by creators

### E) Evaluate Network

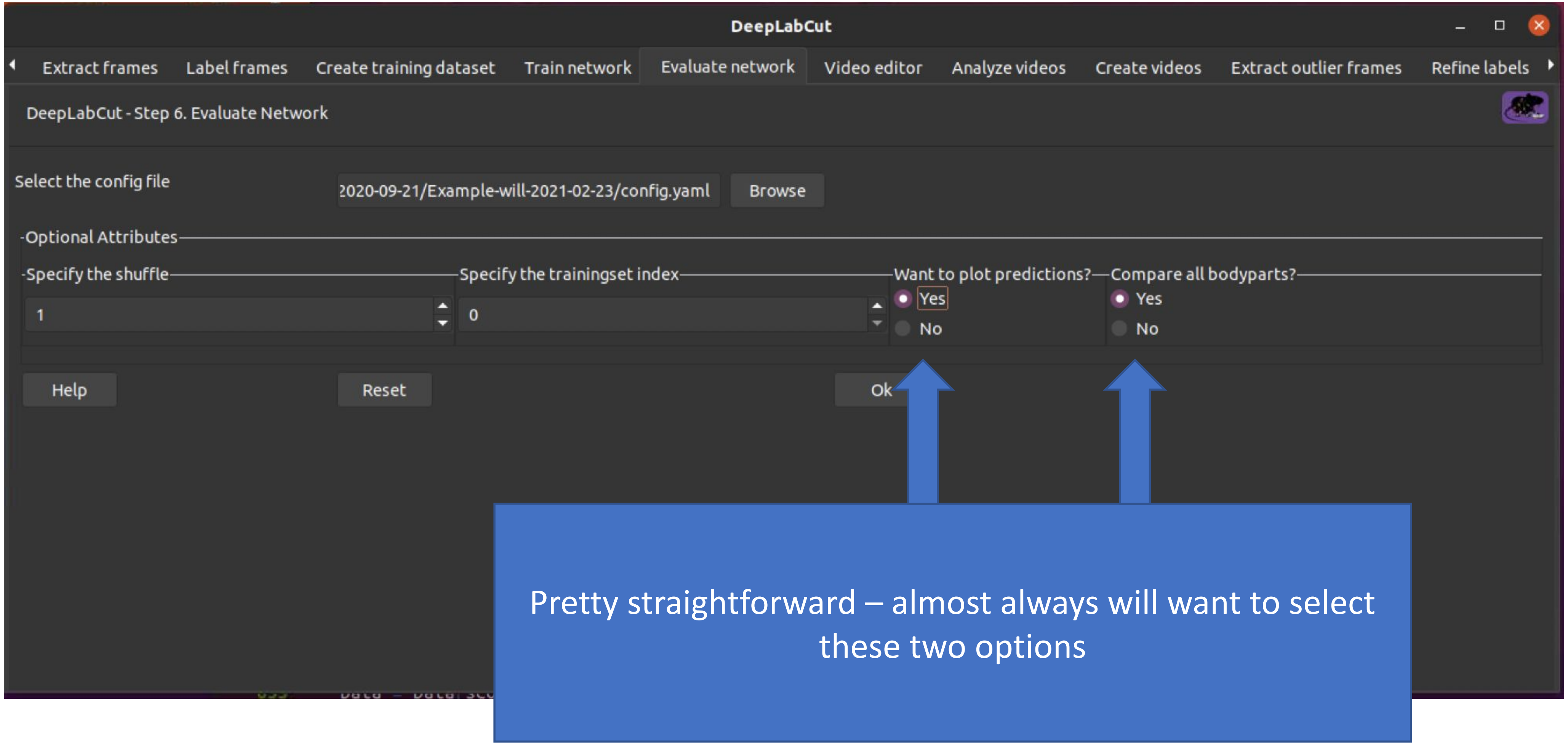

### F) Performing markerless capture using trained model

Just find add videos here

x, y, and certainty values for every marker in every frames

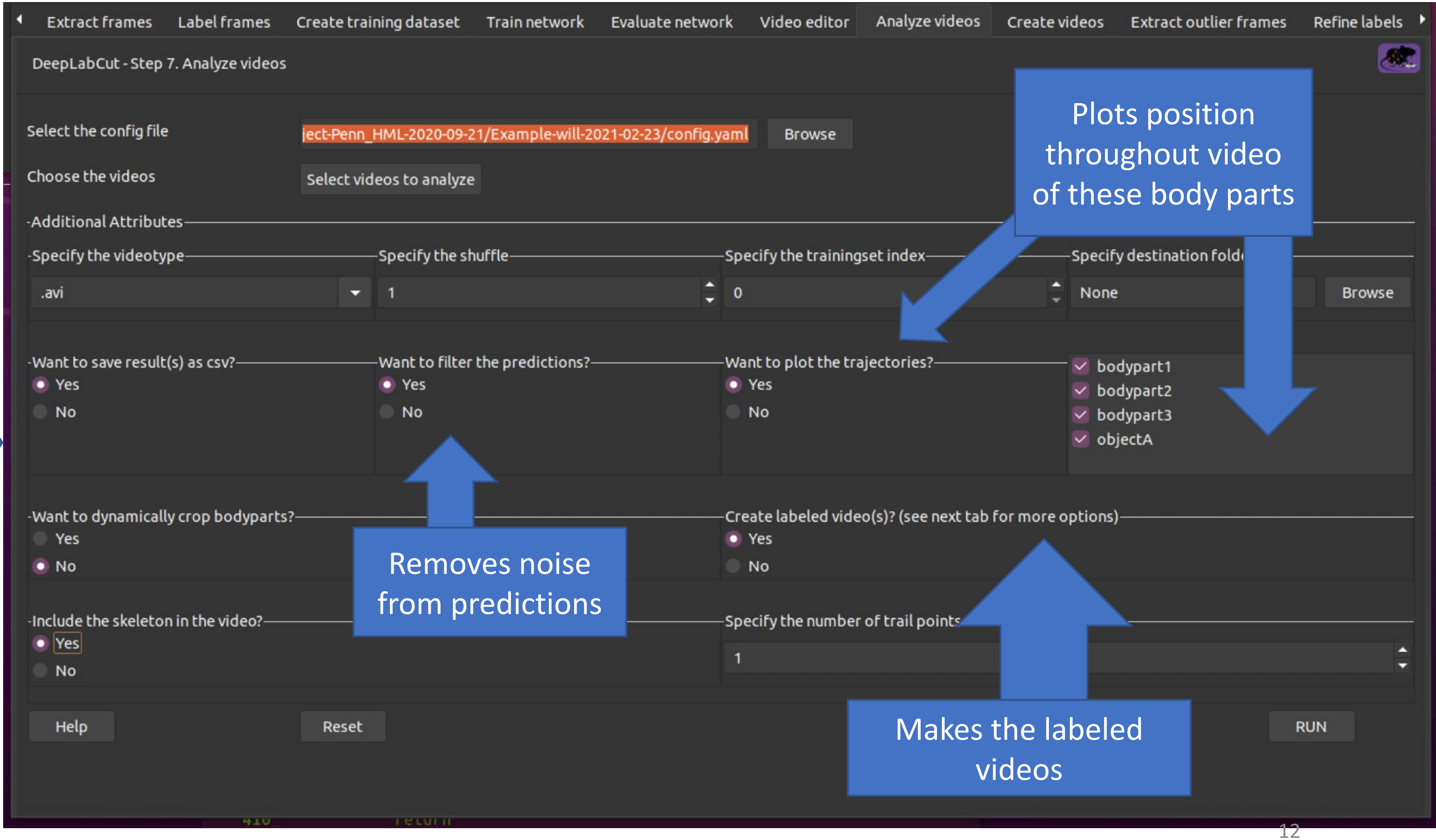


Plots position throughout video of these body parts

Removes noise from predictions

Makes the labeled videos

### Some final notes:

- For installing DeepLabCut and suggested system requirements, the Optogenetics and Neural Engineering Core at Colorado School of Medicine has a useful guide.
- The models produced using this process are heavily influenced by the properties/characteristics of the images used to train the model.
- Some properties that can influence model performance:
  - Lighting
  - Camera quality and position
  - Clothes and other subject characteristics
- Generally, try to train your model using training images that are as close as possible to your anticipated use case.
